# Pivotal involvement of the CX3CL1-CX3CR1 axis for the recruitment of M2-TAMs in skin carcinogenesis

**DOI:** 10.1101/700914

**Authors:** Yuko Ishida, Yumi Kuninaka, Yuki Yamamoto, Mizuho Nosaka, Akihiko Kimura, Fukumi Furukawa, Naofumi Mukaida, Toshikazu Kondo

## Abstract

We previously revealed the crucial roles of CX3CL1-CX3CR1 axis in skin wound healing. Although repeated wounds frequently develop into skin cancer, the roles of CX3CL1 in skin carcinogenesis remain elusive. Here, we proved that CX3CL1 protein expression and CX3CR1^+^ macrophages were observed in human skin cancer tissues. Similarly, we observed the enhancement of CX3CL1 expression and the abundant accumulation of CX3CR1^+^ tumor-associated macrophages (TAMs) with M2 phenotypes in the skin carcinogenesis process induced by the combined treatment with 7,12-dimethylbenz-(a)anthracene (DMBA) and 12-*O*-tetradecanoylphorbol-13-acetate (TPA). In this mouse skin carcinogenesis process, CX3CR1^+^ TAMs exhibited M2 phenotypes with the expression of Wnt3a and angiogenic molecules including vascular endothelial growth factor (VEGF) and matrix metalloproteinase (MMP)-9. Compared to wild-type mice, CX3CR1-deficient mice showed fewer numbers of skin tumors with a lower incidence. Concomitantly, M2-macrophage numbers and neovascularization reduced with the depressed expression of angiogenic factors and Wnt3a. Thus, the CX3CL1-CX3CR1 axis can crucially contribute to skin carcinogenesis by regulating the accumulation and functions of TAMs. Thus, this axis can be a good target for preventing and/or treating skin cancers.

## INTRODUCTION

Inflammation is a classical and biological response to either internal or external stimuli to prevent tissue damages. However, prolonged inflammation destroys the tissue parenchyma, occasionally resulting in the induction of tumorigenesis, which can be furthered by infiltrating leukocytes, which are essential cell components of inflammation (Mantovani *et al*, 2008; Coussens & Werb, 2002).

Chemokines are a family of cytokines, comprising more than 40 molecules: they are produced by various types of cells in response to inflammatory cytokines, growth factors, and pathogenic stimuli (Rossi & Zlotnik, 2000). Chemokines can induce the migration of distinct types of leukocytes and non-leukocytic cells, which express their cognate receptors. As a result, they can crucially contribute to inflammation by inducing the migration of distinct sets of leukocytes into inflammatory sites. Moreover, accumulating evidence may indicate that several chemokines produced by cancer cells can regulate leukocyte trafficking into the tumor microenvironment (Balkwill, 2012; Balkwill, 2004; Allavena *et al*, 2011). Recruited leukocytes can induce angiogenesis and the generation of the fibroblast stroma, which are essential components of cancer tissues (Allavena *et al*, 2011; Mantovani *et al*, 2010), and can eventually promote carcinogenesis. Furthermore, some chemokines can have direct effects on cancer cells, enhancing their growth and motility, thereby accelerating cancer progression (Balkwill, 2004).

CX3C chemokine ligand-1 (CX3CL1) is a membrane-bound chemokine, in contrast to most other chemokines, which are secreted molecules. CX3CL1 can specifically and exclusively bind its receptor, CX3C chemokine receptor-1 (CX3CR1), which is expressed mainly by macrophages, Th1 cells, NK cells, immature dendritic cells, and endothelial cells (Sugaya, 2015). Thus, the CX3CL1-CX3CR1 axis can be used to augment tumor immunity by recruiting Th1 and NK cells (Sugaya, 2015). In contrast to its potential anti-tumor activity, accumulating evidence may indicate the pro-tumorigenic activity of the CX3CL1-CX3CR1 axis, which was overexpressed in various types of cancers including breast, prostate, pancreas, ovary, kidney, and colon cancers (Tsang *et al*, 2013; Shulby *et al*, 2004; Celesti *et al*, 2013; Kim *et al*, 2012; Yao *et al*, 2014; Zheng *et al*, 2013). Similarly, the roles of CX3CL1 in tumor invasion and metastasis are still controversial. CX3CL1 can promote cancer metastasis using various routes including bloodstream, lymphatic vessels, and nerves (Borsig *et al*, 2014), whereas CX3CL1 can prevent glioma invasion by promoting tumor cell aggregation and eventually reducing their invasiveness (Sciumè *et al*, 2010). Thus, the roles of the CX3CL1-CX3CR1 axis in carcinogenesis processes remain elusive.

We have previously demonstrated that the interaction between locally produced CX3CL1 and CX3R1-expressing cells contributed crucially to the skin wound healing process by promoting the macrophage accumulation and functions of macrophages (Ishida *et al*, 2008). Based on similar changes common to the wound healing process and tumorigenesis, Dvorak proposed that tumors are wounds that do not heal (Dvorak, 2015). This proposition prompted us to investigate the roles of the CX3CL1-CX3CR1 axis in skin carcinogenesis. We demonstrated that CX3CL1 protein expression and CX3CR1-expressing macrophages were observed in human skin cancer tissues. In order to address the pathogenic roles of the CX3CL1-CX3CR1 axis in skin carcinogenesis, we utilized 7,12-dimethylbenz-(a)anthracene (DMBA)/12-*O*-tetradecanoylphorbol-13-acetate (TPA)-induced two-stage skin carcinogenesis, which mimics the multistage development of human skin cancer (Kemp, 2005). Here, we provided definitive evidence to indicate the vital roles of the CX3CL1-CX3CR1 axis in this skin carcinogenesis model.

## RESULTS

### CX3CL1 and CX3CR1 expression in human skin cancer tissues of basal cell carcinoma (BCC) and squamous cell carcinoma (SCC)

The importance of the interaction between CX3CL1 and CX3CR1 has been implicated in many inflammatory diseases, including rheumatoid arthritis, vasculitis, systemic sclerosis, and atopic dermatitis (Echigo *et al*, 2004; Hasegawa *et al*, 2005; Morimura *et al*, 2013; Murphy *et al*, 2008). Hence, initially, we immunohistochemically examined the expression of CX3CL1 and CX3CR1 in the BCC and SCC tissues. CX3CL1 (Figure 1A) and CX3CR1 (Figure 1B) were mainly detected in infiltrating leukocytes in all specimens from both the BCC or SCC lesions that we examined. Double-color immunofluorescence analyses demonstrated that CD68^+^ macrophages, but not CD31^+^ endothelial cells, were the main cellular source of CX3CL1 (Figure 1C and Supplemental figure 1A). Moreover, CX3CR1 was detected in both CD68^+^ macrophages and endothelial cells (Figure 1, D and E). Most CX3CL1^+^ cells consistently expressed CX3CR1 (Supplemental Figure 1B). These observations indicate the potential involvement of the CX3CL1-CX3CR1 axis in skin carcinogenesis, probably via its action on tumor-associated macrophages (TAMs), which simultaneously express CX3CL1 and CX3CR1.

**Figure 1.**
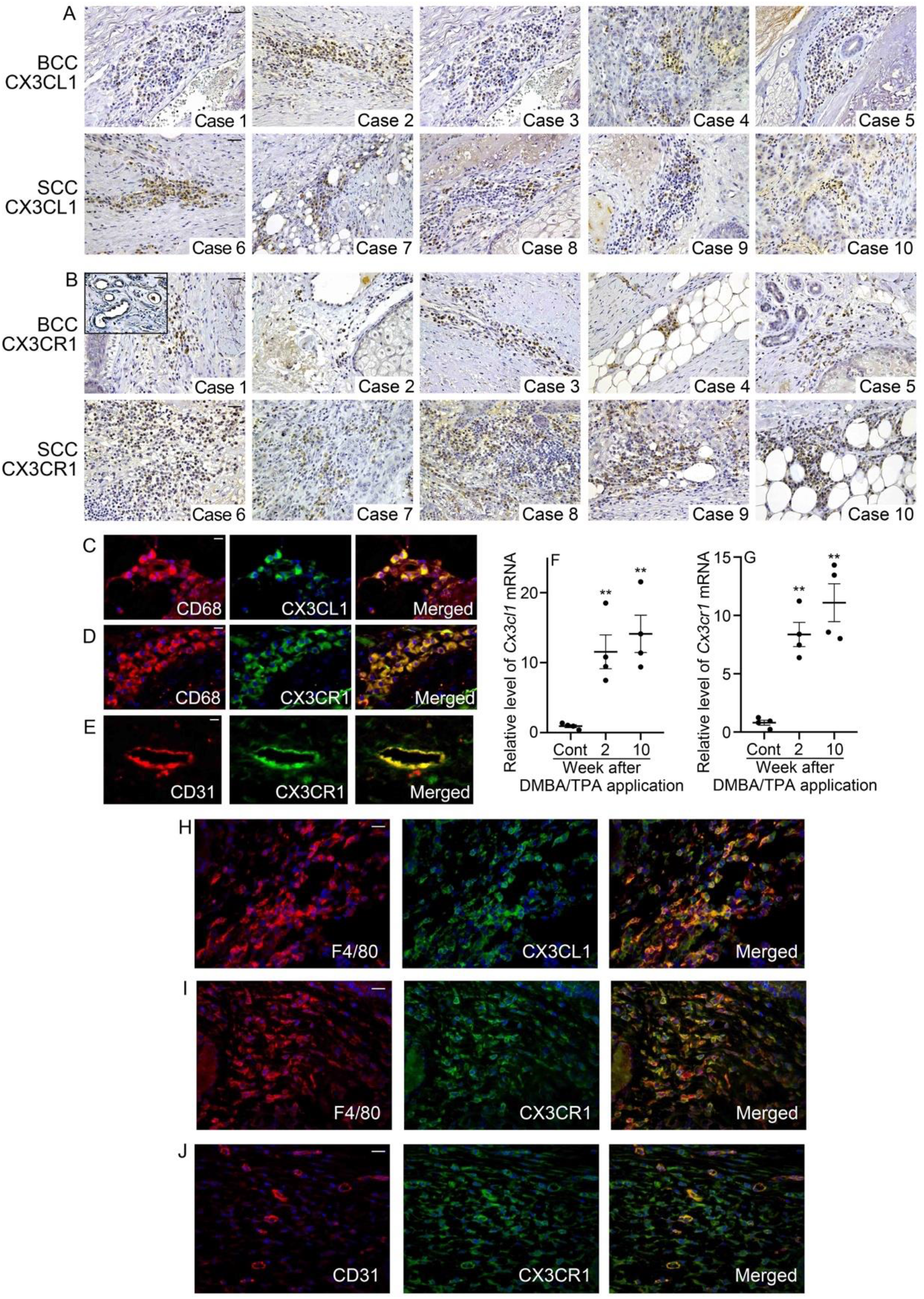
CX3CL1 and CX3CR1 expression in skin cancer. (A and B) The expression of CX3CL1 (A) and CX3CR1 (B) in the human BCC (Case 1-5) and SCC tissues (Case 6-10). The samples were processed to immunohistochemical analysis using anti-CX3CL1 or anti-CX3CR1 antibodies. Representative results are shown here. Scale bars, 50 μm; scale bars in inserts, 40 μm. (C-E) Cell types expressing CX3CL1 and CX3CR1 in the human skin cancer. Double-color immunofluorescence analyses were performed on human skin cancer tissues. Representative results are shown here. Signals were merged digitally. Scale bars, 20 μm. (F and G) The expression of *Cx3cl1* (F) and *Cx3cr1* (G) mRNA in the skin of WT mice after DMBA/TPA treatment. Quantitative RT-PCR analyses of *Cx3cl1* and *Cx3cr1* mRNA was carried out. Values represent mean ± SEM (n=6). *, *P* < 0.05; **, *P* < 0.01, vs. unchallenged skin, by 2-way ANOVA followed by Dunnett’s post-hoc test. (H-J) Cell types expressing CX3CL1 and CX3CR1 in the skin of DMBA/TPA-treated WT mice. Double-color immunofluorescence analyses were performed. Representative results from six individual animals are shown here. Signals were merged digitally. Scale bars, 20 μm.

### CX3CL1 and CX3CR1 expression in the skin after DMBA/TPA treatment

Next, we examined the *Cx3cl1* and *Cx3cr1* mRNA expression in the skin of WT mice after DMBA/TPA treatment. Both *Cx3cl1* and *Cx3cr1* mRNAs were faintly detected in the skin of the untreated WT mice. DMBA/TPA treatment significantly enhanced the *Cx3cl1* and *Cx3cr1* mRNA expression in the skin later than two weeks after DMBA application (Figure 1F and G). The double-color immunofluorescence analyses further demonstrated that F4/80^+^ cells were the major cellular source of CX3CL1 (Fig. 1H), similar to the observation of the human skin cancer tissues. CX3CR1 protein was mainly detected in F4/80^+^ macrophages (Figure 1I), and to a lesser degree in endothelial cells (Figure 1J), with is consistent with our data of the human skin cancer tissues. Thus, CX3CL1 and CX3CR1 expression was enhanced in chemical-induced mouse skin carcinogenesis, as well as in human skin cancer tissues.

### The pathogenic involvement of the CX3CR1-CX3CL1 axis in skin carcinogenesis

In order to address the roles of the CX3CL1-CX3CR1 axis in skin carcinogenesis, WT and *Cx3cr1*^-/-^ mice were subjected to DMBA/TPA treatment. There were no apparent differences between the skin structures of the unchallenged WT and *Cx3cr1*^-/-^ mice (Figure 2A). On the contrary, a marked epidermal thickness was observed 20 weeks after DMBA/TPA treatment in WT, but not *Cx3cr1*^-/-^ mice (Figure 2, A and B). Consistently, the number of Ki67^+^ proliferating epidermal cells were significantly larger in WT mice than in *Cx3cr1*^-/-^ mice (Figure 2, C and D). Several lines of evidence demonstrated that the activation of EGFR signal pathway was closely involved in skin carcinogenesis (Tardáguila *et al*, 2013; Sibilia *et al*, 2000; Hara *et al*, 2005). Actually, EGFR signals were less activated in *Cx3cr1*^-/-^ mice, compared with WT ones (Figure 2, E and F). Although both strains started to develop papillomas later than 10 weeks after DMBA/TPA treatment, the tumor incidence was higher in WT mice than in *Cx3cr1*^-/-^ mice at all time points that we examined (Figure 2, G and H). As a consequence, at 20 weeks after DMBA/TPA treatment, 90% of the WT mice, but only 40% of the *Cx3cr1*^-/-^ mice developed papillomas (Figure 2H). Moreover, the average numbers of papillomas in *Cx3cr1*^-/-^ mice were significantly lower than those in WT mice (Figure 2I). These observations would imply that the contribution of the CX3CL1-CX3CR1 axis may contribute to chemical-induced skin carcinogenesis by inducing the proliferation of epidermal cells. Next, we explored the contribution of BM-derived CX3CR1^+^ cells to skin carcinogenesis by using BM chimeric mice generated from WT and *Cx3cr1*^-/-^ mice. Both WT and *Cx3cr1*^-/-^ mice transplanted with WT mouse-derived BM cells exhibited a higher tumor incidence than the recipients of *Cx3cr1*^-/-^ mouse-derived BM cells (Figure 2J). The average numbers of papillomas in the mice receiving *Cx3cr1*^-/-^-BM cells were significantly lower than those in the mice receiving WT-BM cells (Figure 2K). These observations would imply the crucial involvement of radiosensitive BM-derived CX3CR1^+^ cells but not radio-resistant keratinocytes in skin carcinogenesis.

**Figure 2.**
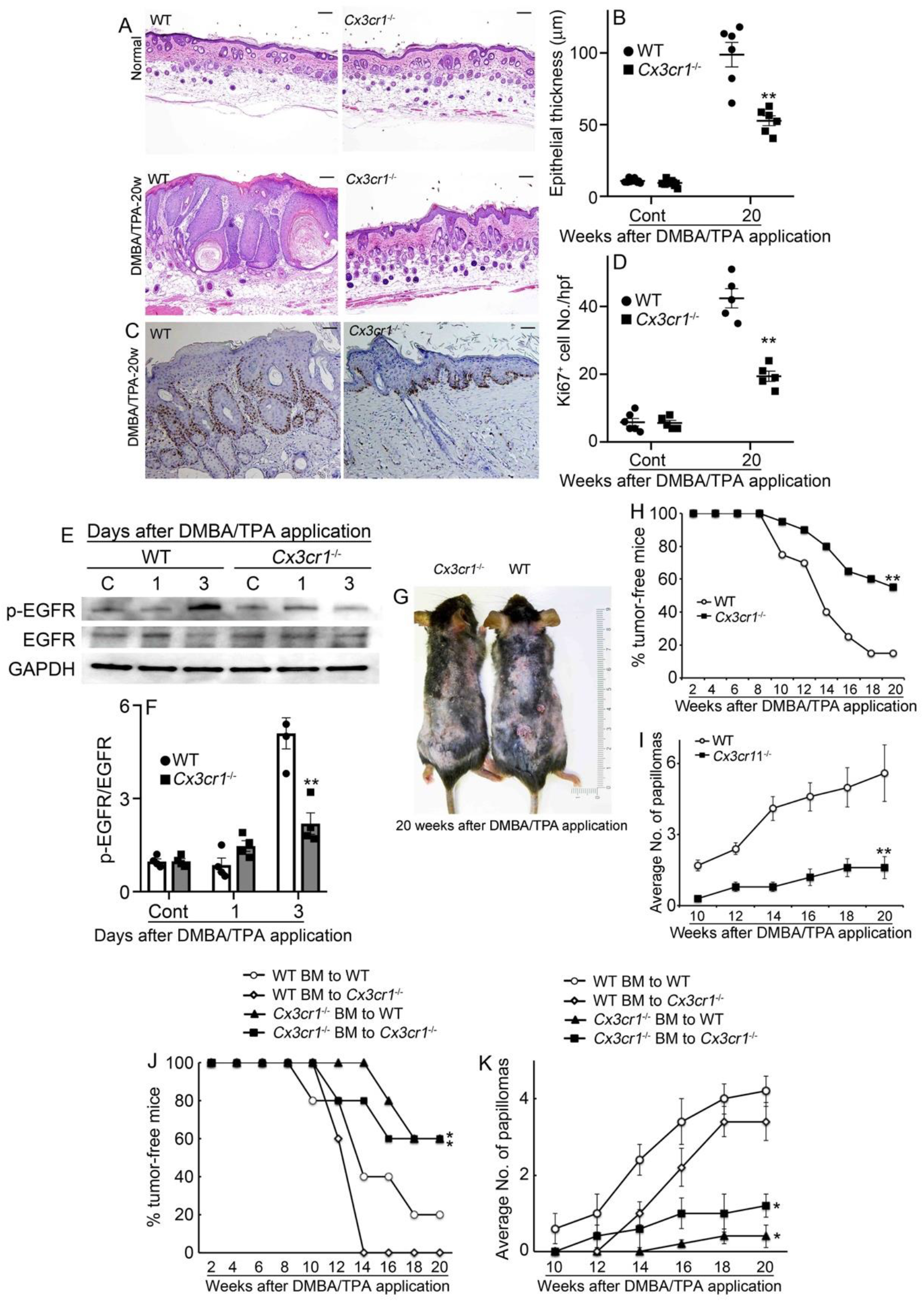
The evaluation of tumor incidence in WT and *Cx3cr1^-/-^* mice after DMBA/TPA treatment. (A) Evaluation of epidermal thickness in non-tumor skin lesion. Histological observations were conducted on the epidermal layers in WT and *Cx3cr1^-/-^* mice before or after DMBA/TPA treatment. Representative results from six animals are shown here. Scale bars, 100 μm (HE stain). (B) The average of the epidermal thickness was calculated. All values represent mean ± SEM (n=6). *, *P* < 0.05, WT vs. *Cx3cr1^-/-^* mice, by unpaired Student’s *t* test. (C) Evaluation of epidermal proliferation in non-tumor skin lesion. Ki67 was immunohistochemically detected. Representative results from six animals are shown here. Scale bars, 100 μm. (D) The number of Ki67^+^ cells was counted. All values represent mean ± SEM (n=6). *, *P* < 0.05, WT vs. *Cx3cr1^-/-^* mice, by unpaired Student’s *t* test. (E and F) The effects of CX3CR1 on phosphorylation of EGFR in skin tissue after DMBA/TPA treatment. (E) Western blotting analysis using anti-GAPDH antibody confirmed that an equal amount of protein was loaded onto each lane. (F) The ratio of p-EGFR/EGFR was densitometrically determined and are shown. All values represent means ± SEM (4 independent experiments). \*\**P*<0.01 vs. WT mice. (G) Macroscopic pictures of skin papillomas in WT and *Cx3cr1^-/-^* mice. Representative results at 20 weeks after DMBA/TPA treatment are shown here. (H) The percentage of tumor-free mice at the indicated time points after DMBA/TPA treatment (n=20). **, *P* < 0.01, WT vs. *Cx3cr1^-/-^* mice. (I) The average number of skin papillomas per mouse at the indicated time points after DMBA/TPA treatment (n=20). **, *P* < 0.01, WT vs. *Cx3cr1^-/-^* mice. (J) The percentage of tumor-free mice at the indicated time points after DMBA/TPA treatment (n=5). *, *P* < 0.05, vs. WT-BM to WT. (K) The average number of skin papillomas per mouse at the indicated time points after DMBA/TPA treatment (n=5). *, *P* < 0.05, vs. WT-BM to WT, by unpaired Student’s *t* test.

### The effects of CX3CR1 deficiency on macrophage recruitment after DMBA/TPA treatment

CX3CR1 expression in TAMs prompted us to examine the effects of CX3CR1 deficiency on macrophage infiltration in this carcinogenesis process. At one week after DMBA/TPA treatment, the recruitment of F4/80^+^ macrophages, but not Ly-6G^+^CD11b^+^ neutrophils and CD3^+^ lymphocytes, was significantly reduced in *Cx3cr1*^-/-^ mice, compared to the case in WT mice (Figure 3, A and B). Moreover, even at 10 weeks after DMBA/TPA treatment, intradermal macrophage recruitment was attenuated in *Cx3cr1*^-/-^ mice, compared to the case in WT mice (Figure 3, C and D). Given the important role of macrophage polarization to M2-like phenotype in carcinogenesis (Tariq *et al*, 2017; Isidro & Appleyard, 2016), we determined the expression of M2-related molecules in dermal macrophages isolated from DMBA/TPA-treated WT or *Cx3cr1*^-/-^ mice. The numbers of the isolated whole macrophages were significantly reduced in *Cx3cr1*^-/-^ mice, compared to the WT mice, both at 48 and 72 h after DMBA/TPA treatment (Figure 3E). Moreover, the *Cx3cr1*^-/-^-derived dermal macrophages exhibited a reduced expression of M2-macrophage markers such as *Mrc1/Cd206, Cd163, Il10, Ccl17*, and *arginase1*, compared to the WT-derived dermal macrophages (Figure 4, F-J). Consistently, even at five days after DMBA/TPA treatment, CD68^+^CD206^+^ M2-like macrophages numbers were reduced in *Cx3cr1*^-/-^ mice, compared to the case in WT mice (Figure 3, K and L). Furthermore, nearly three-quarters of the CX3CR1^+^ macrophages expressed CD206, an M2-macrophage marker (Figure 3M). A triple-color immunofluorescence analysis demonstrated that a large number of F4/80^+^CD206^+^ M2 macrophages expressed CX3CR1 in DMBA/TPA-induced carcinogenesis (Figure 3N). Consistently, the expression of CD163, a human M2 marker, was conspicuous in human skin cancer lesions, particularly SCC ones, compared with that of HLA class II, an M1 marker (Figure 4, A-C). A triple-color immunofluorescence analysis demonstrated that a large number of CD68^+^CD163^+^ M2 macrophages expressed CX3CR1 in human SCC lesions (Figure 4D). Thus, M2 macrophage infiltration could be a prominent feature of skin carcinogenesis in mice and humans. Finally, DMBA/TPA-treated WT and *Cx3cr1*^-/-^ mice displayed similar levels of gene expression of the *Ccl2* and *Ccl3* genes, which exhibit potent chemotactic activities for macrophages but utilize distinct receptors than CX3CR1 (Supplemental figure 2). These observations imply that DMBA/TPA treatment induced the infiltration of macrophages and their subsequent polarization into the M2 phenotype by inducing CX3CL1, which can act on the CX3CR1 expressed on macrophages.

**Figure 3.**
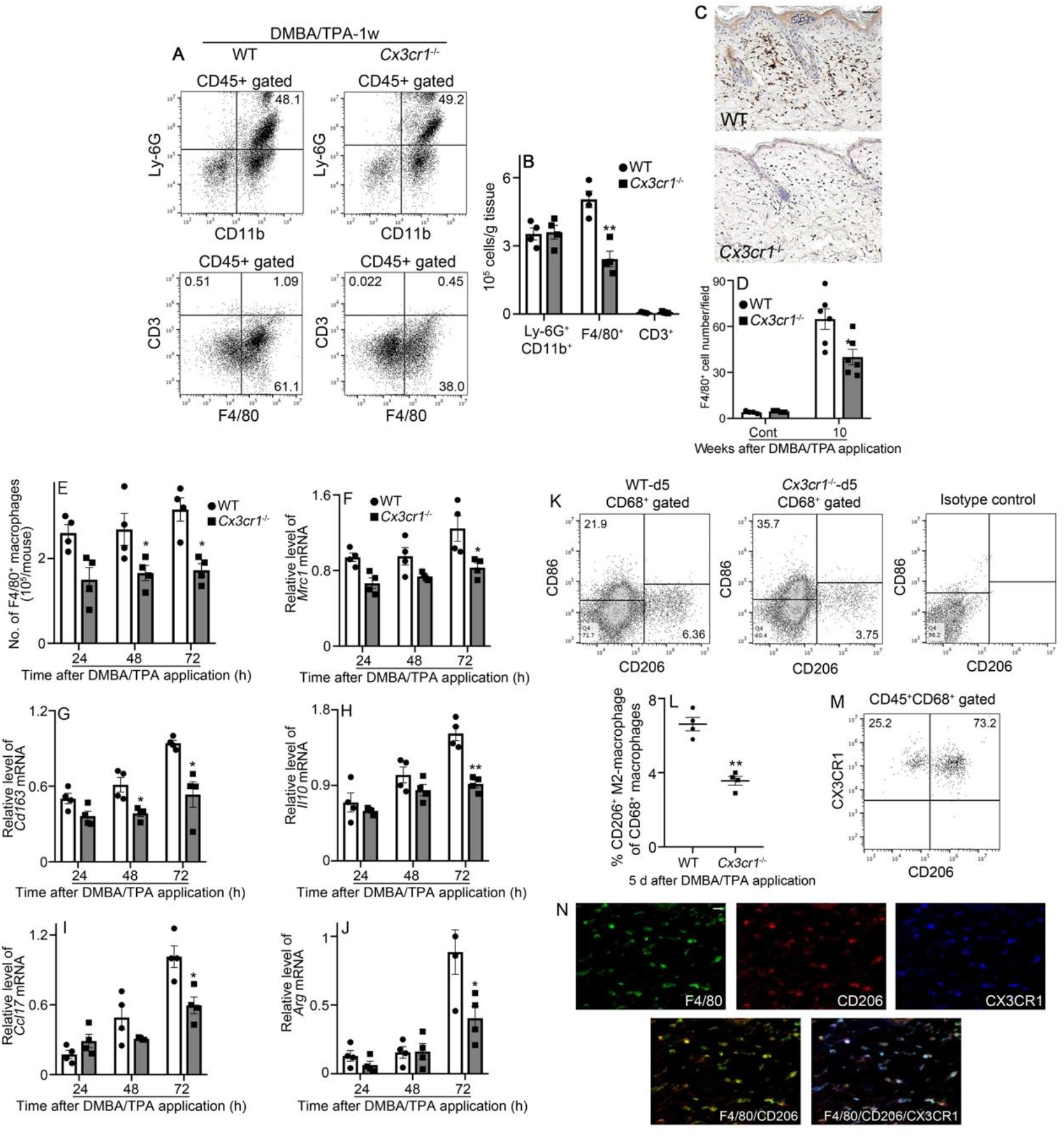
Evaluation of leukocyte recruitment in WT and *Cx3cr1^-/-^* mice after DMBA/TPA treatment. (A) Flow cytometric analysis on CD45^+^ leukocytes in the skin tissue of WT and *Cx3cr1*^-/-^ mice at 1 week after DMBA/TPA treatment. Representative results from 6 independent experiments are shown. (B) The number of Ly-6G^+^CD11b^+^ neutrophils, F4/80^+^ macrophages, and CD3^+^ lymphocytes were calculated, and are shown. Each value represents mean ± SEM. \*\**P* < 0.05, vs. WT, by unpaired Student’s *t* test. (C) Immunohistochemical analyses were conducted by using anti-F4/80 mAb. Representative results from six individual animals are shown. Scale bar, 50 μm. (D) The number of macrophages in the skin tissue at 10 weeks after DMBA/TPA treatment was determined. All values represent mean ± SEM (n=6). *, *P* < 0.05, WT vs. *Cx3cr1^-/-^* mice, by unpaired Student’s *t* test. (E-J) Suppressed gene expression of M2-macrophage markers in tissue macrophages obtained from *Cx3cr1*^-/-^ mice compared to those of WT mice. (E) Cell number of F4/80^+^ macrophages in skin tissues of WT and *Cx3cr1*^-/-^ mice at 24, 48, and 72 h after DMBA/TPA treatment. F4/80^+^ cells were extracted from skin single cell suspension by using MACS system, and calculated as cell number per mouse. Values represent mean ±SEM (n=4). \**P* < 0.05, vs. WT. (F-J) The expression of M2-macropahge marker, *Mrc1* (F), *Cd163* (G), *Il10* (H), *Ccl17* (I), and *Arg* (J) mRNA in the skin of WT mice after DMBA/TPA treatment. Values represent mean ± SEM (n=4). *, *P* < 0.05; **, *P* < 0.01, vs. WT, by 2-way ANOVA followed by Dunnett’s post-hoc test. (K-N) Depressed M2-macrophage numbers of skin tumor sites of *Cx3cr1*^-/-^ mice, compared with those of WT mice. (K) Flow cytometric analysis on CD86^+^ M1-macrophages and CD206^+^ M2-macrophages in the skin tissue of WT and *Cx3cr1*^-/-^ mice at 5 days after DMBA/TPA treatment. Representative results from 6 independent experiments are shown. (L) Percentage of CD206^+^ M2-macrophage among CD68^+^ macrophages (n=4). (M) Flow cytometric analysis on the portion of CD206^+^ M2-macrophages co-expressing CX3CR1. Representative results experiments are shown. (N) Triple-color immunofluorescence analyses were performed on DMBA/TPA-treated tissues (10 w). Representative results are shown here. Signals were merged digitally. Scale bar, 20 μm.

**Figure 4.**
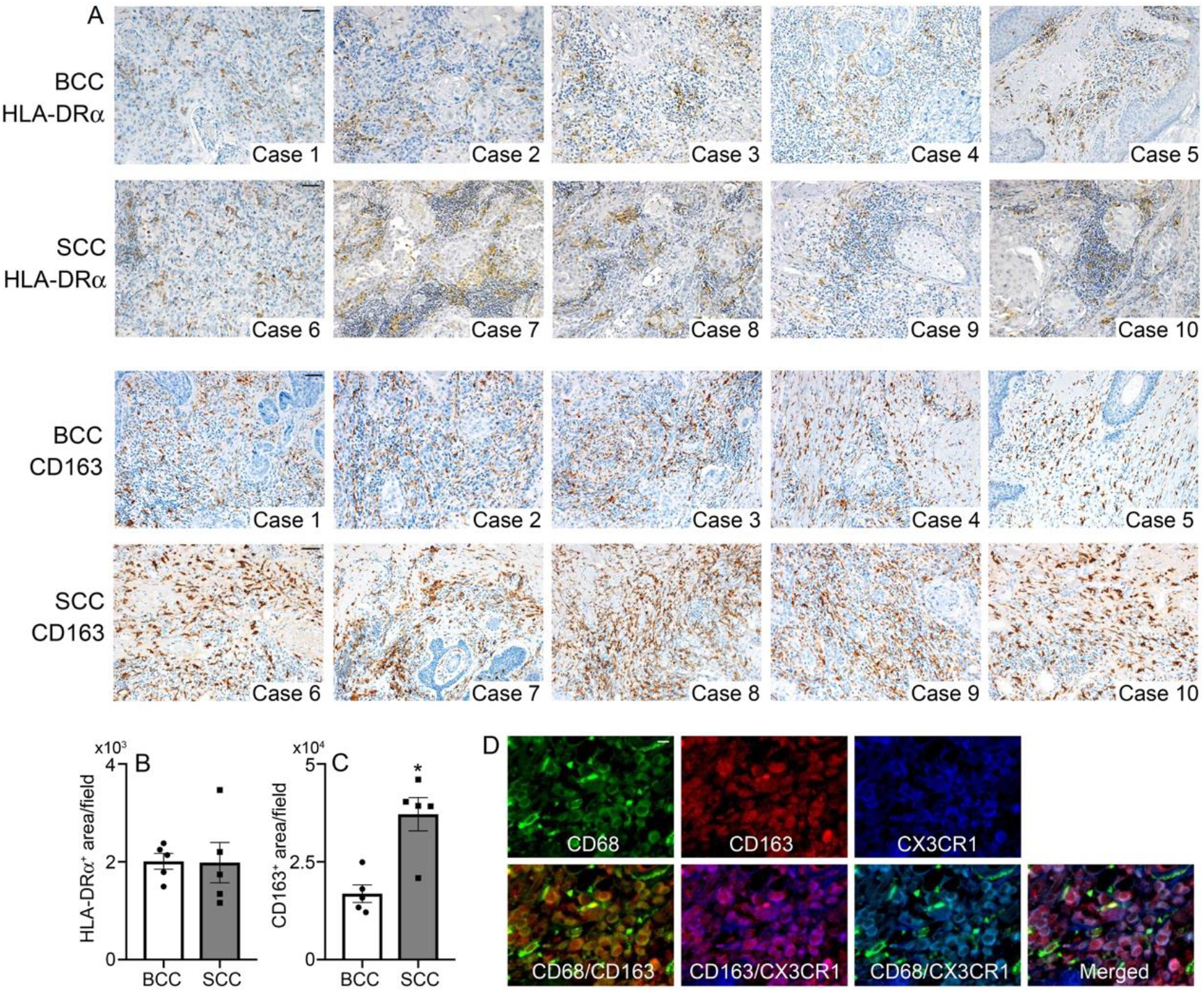
M1 and M2 macrophages in human skin cancer. (A) The expression of HLA-DRα (M1 macrophages marker) and CD163 (M2 macrophages marker) in the human BCC (Case 1-5) and SCC tissues (Case 6-10). Representative results are shown here. Scale bars, 50 μm. (B and C) The occupied degrees of HLA-DRα^+^ area (B) or CD163^+^ area (C) per high power field in the skin tissue were determined. All values represent mean ± SEM (n=5). \**P* < 0.05, BCC, by unpaired Student’s *t* test. (D) CX3CR1-expressing M2 macrophages in human SCC tissues. Triple-color immunofluorescence analyses were performed on human SCC tissues. Representative results are shown here. Signals were merged digitally. Scale bar, 20 μm.

### Reduced neovascularization in mice lacking CX3CR1 after DMBA/TPA treatment

CX3CR1 expression by CD31-positive cells incited us to examine the effects of CX3CR1 deficiency on angiogenesis during the course of skin carcinogenesis. DMBA/TPA treatment increased the intradermal vessel density in WT mice, but these increments were significantly reduced in *Cx3cr1*^-/-^ mice (Figure 5, A and B). Moreover, WT mice exhibited enhanced intradermal gene expression of *Vegf* and *Mmp9*, two potent angiogenic molecules, compared to the case in *Cx3cr1*^-/-^ mice (Figure 5, C and D). Moreover, the VEGF and MMP9 proteins were detected mainly in CX3CR1^+^ macrophages (Figure 5, E and F), suggesting that CX3CR1^+^ M2-like macrophages could contribute to angiogenesis in the development of chemical-induced skin carcinogenesis. Thus, the absence of the CX3CR1 axis reduced neovascularization directly and indirectly by reducing the expression of the potent angiogenic factors, VEGF and MMP9, during the course of DMBA/TPA-induced skin carcinogenesis.

**Figure 5.**
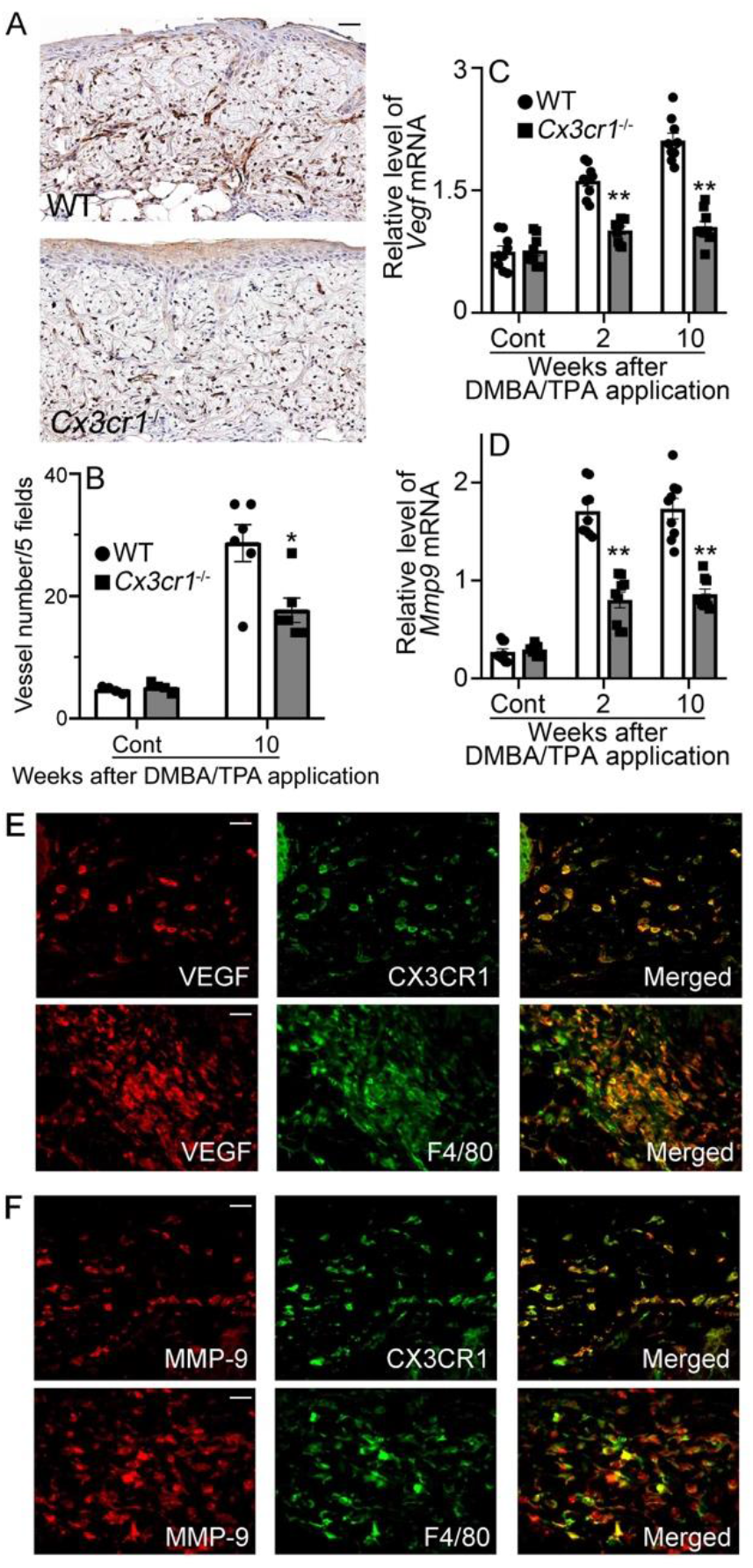
Evaluation of intradermal neovascularization in WT and *Cx3cr1^-/-^* mice after DMBA/TPA treatment. (A) Immunohistochemical analyses were conducted by using anti-CD31 mAb. Representative results from six individual animals are shown here. Scale bar, 50 μm. (B) The numbers of vessels in the skin tissue were determined. All values represent mean ± SEM (n=6). *, *P* < 0.05, WT vs. *Cx3cr1^-/-^* mice, by unpaired Student’s *t* test. (C) Intradermal gene expression of *Vegf* in WT and *Cx3cr1*^-/-^ mice after DMBA/TPA treatment. Quantitative RT-PCR analyses were carried out. Values represent mean ± SEM (n=6). (D) Intradermal gene expression of *Mmp9* in WT and *Cx3cr1*^-/-^ mice after DMBA/TPA treatment. Values represent mean ± SEM (n=6). (E) Cell types expressing VEGF in the skin of DMBA/TPA-treated WT mice. (F) Cell types expressing MMP-9 in the skin of DMBA/TPA-treated WT mice. Double-color immunofluorescence analyses were performed. Representative results from six individual animals are shown here. Signals were merged digitally. Scale bars, 20 μm.

### Reduced expression of Wnt3a in DMBA/TPA-treated CX3CR1**^-/-^** mice

The crucial roles of β-catenin-mediated signaling in a variety of skin cancers (Gat *et al*, 1998; Malanchi *et al*, 2008) provoked us to determine the gene expression of *Wnt3a*. Indeed, DMBA/TPA treatment enhanced the intradermal *Wnt3a* mRNA expression in WT mice later than six weeks after the treatment, but this enhancement was significantly attenuated in *Cx3cr1*^-/-^ mice (Figure 6A). Wnt3a^+^ cells were consistently detected in the skin of the WT and *Cx3cr1*^-/-^ mice, at 10 weeks after the DMBA/TPA treatment (Figure 6B), but the number of Wnt3a^+^ cells was significantly lower in *Cx3cr1*^-/-^ mice than in WT mice (Figure 6C). A double-color immunofluorescence analysis detected Wnt3a in CX3CR1^+^ and F4/80^+^ cells, indicating that CX3CR1^+^ M2-like macrophages could produce Wnt3a, eventually contributing to chemical-induced skin carcinogenesis (Figure 6, D and E). Moreover, CX3CL1 augmented the Wnt3a expression in a mouse macrophage cell line, RAW264.7 (Figure 6F). Immunohistochemical analysis demonstrated that the nuclear accumulation of β-catenin in the epidermal cells was markedly decreased in *Cx3cr1*^-/-^ mice, compared to the case in WT mice, at 10 weeks after DMBA/TPA treatment (Figure 6G). The amount of unphosphorylated (active) β-catenin protein was increased in WT mice at 10 weeks after DMBA/TPA treatment but this increase was significantly lowered in *Cx3cr1*^-/-^ mice (Figure 6, H and I). Moreover, DMBA/TPA treatment increased the *Cox2* expression in the skin of WT mice, but these increments were significantly lowered in *Cx3cr1*^-/-^ mice at two weeks after DMBA/TPA treatment (Figure 6J). A double-color immunofluorescence analysis detected COX-2 in F4/80^+^ and CX3CR1^+^ cells (Figure 6, K and L). Several lines of evidence have indicated that COX-2/prostaglandin E2 can exert pro-oncogenic actions through β-catenin signaling (Castellone *et al*, 2005; Shao *et al*, 2005). Thus, locally produced CX3CL1 can recruit Wnt3a- and COX-2-expressing macrophages, and can further enhance their Wnt3a expression, thereby activating the β-catenin signaling pathway and eventually skin carcinogenesis.

**Figure 6.**
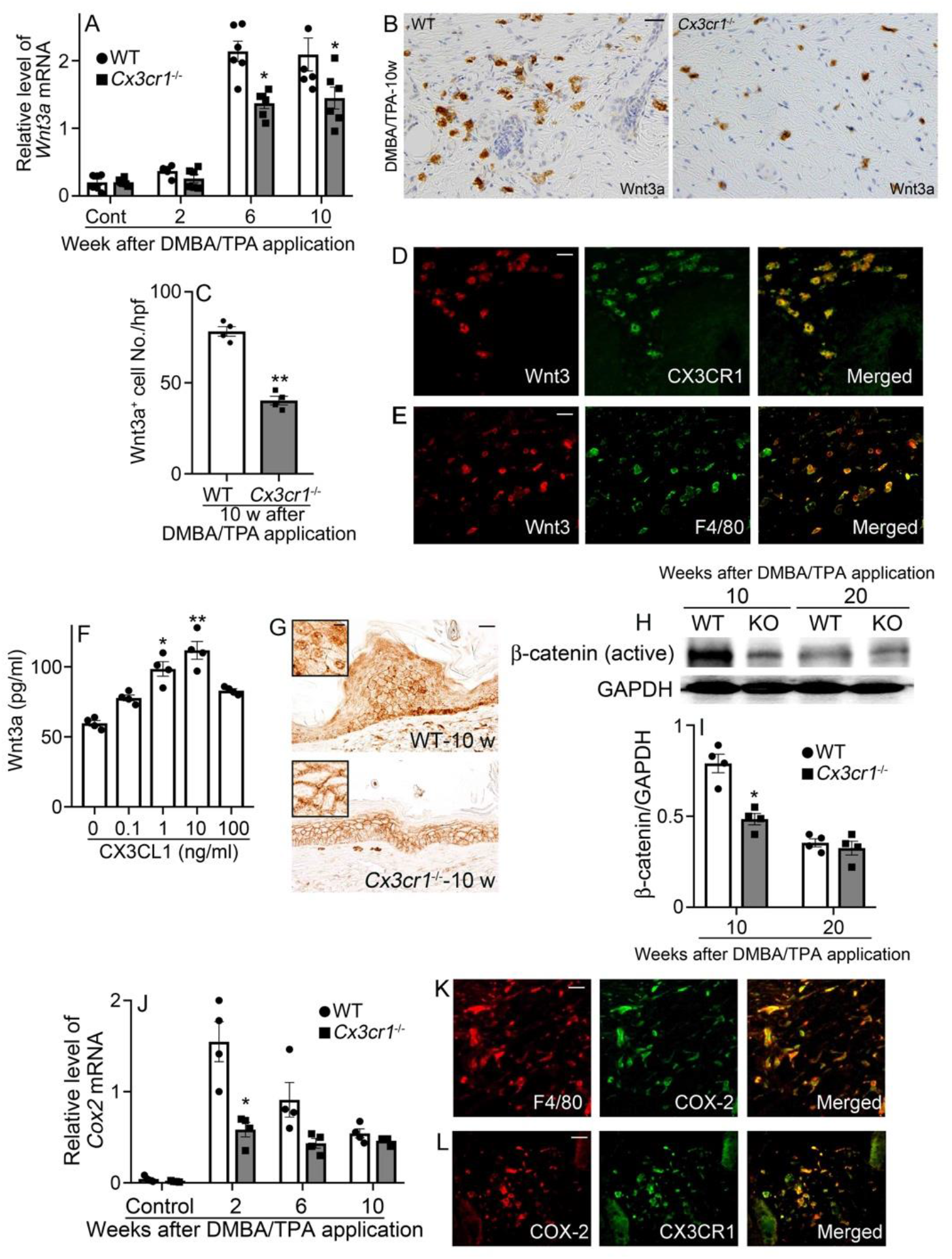
Evaluation of intradermal Wnt3a expression in WT and *Cx3cr1*^-/-^ mice after DMBA/TPA treatment. (A) Intradermal mRNA expression of *Wnt3a* in WT and *Cx3cr1*^-/-^ mice after DMBA/TPA treatment. Values represent mean ± SEM (n=6). *, *P* < 0.05, WT vs. *Cx3cr1^-/-^* mice, by 2-way ANOVA followed by Dunnett’s post-hoc test. (B) Immunohistochemical analyses were conducted by using anti-Wnt3a antibodies. Representative results from six individual animals are shown here. Scale bar, 20 μm. (C) The number of Wnt3a^+^ cells in the skin tissue were determined. All values represent mean ± SEM (n=6). *, *P* < 0.05, WT vs. *Cx3cr1^-/-^* mice, by unpaired Student’s *t* test. (D and E) Cell type expressing Wnt3 in the skin of DMBA/TPA-treated WT mice. Representative results from six individual animals are shown here. Signals were merged digitally. Scale bars, 20 μm. (F) RAW263.7 were stimulated with CX3CL1 for 2 h to be subjected to ELISA for Wnt3a. All values represent the mean ± SEM (n=4 independent experiments). \*\**P*<0.01, \**P*<0.05, vs. no stimulation, by one-way ANOVA followed by Dunnett’s post hoc test. (G) Skin samples were immunostained with anti-β-catenin antibodies and representative result from 4 independent animals were shown. Insets are higher magnifications of the positively stained cells. Scale bar, 20 μm; scale bar in inserts, 5 μm. (H) The expression of β-catenin (active) and GAPDH were analyzed by Western blot analysis. Representative image from 4 independent experiments are shown. (I) β-catenin (active) protein were determined based on the band intensities. All values represent the mean ± SEM. \**P* < 0.05, vs. WT, by unpaired Student’s *t* test. (J) Intradermal gene expression of *Cox2* in WT and *Cx3cr1*^-/-^ mice after DMBA/TPA treatment. Value represent mean ± SEM (n=4). \**P* < 0.05, vs. WT, by 2-way ANOVA followed by Dunnett’s post-hoc test. (K and L) Cell types expressing COX-2 in the skin of DMBA/TPA-treated WT mice. Representative results from six individual animals are shown here. Signals were merged digitally. Scale bars, 20 μm.

## DISCUSSION

The crucial involvement of the CXCL1-CX3CR1 axis in skin wound healing incited us to examine the CX3CL1 and CX3CR1 expression in human skin cancer tissues. We detected abundant CX3CL1 expression in conditions of CX3CR1^+^ cell infiltration in human skin cancer tissues. In order to investigate its role in skin carcinogenesis, we utilized DMBA/TPA-induced two-step skin carcinogenesis, where a single DMBA application induces DNA mutations in epidermal cells, and subsequent repeated exposure to TPA induces chronic inflammation, to accelerate tumorigenesis (DiGiovanni, 1992). Indeed, similar CX3CL1 and CX3CR1 expression patterns were observed in the skin of mice after DMBA/TPA-treatment. Moreover, given the exclusive use of CX3CR1 by its ligand, CX3CL1, reduced DMBA/TPA-induced skin carcinogenesis in CX3CR1^-/-^ mice indicates the crucial involvement of the CX3CL1-CX3CR1 axis in this carcinogenesis model.

TAMs are frequently a major cell component of cancer tissues and usually lack cytotoxic activity, but exhibit M2-like phenotypes with the expression of angiogenic factors (Mantovani *et al*, 2017). Indeed, CX3CR1-expressing TAMs in the present mouse model exhibited M2-like phenotypes and expressed abundantly expressed two potent angiogenic factors, VEGF and MMP-9. CX3CR1 deficiency reduced macrophage infiltration, with depressed M2-like macrophage numbers, and reduced the expression of VEGF and MMP-9. Although TAMs are presumed to be derived mostly from circulating monocytes that are attracted towards tumor sites by locally produced chemokines including CCL2 and CCL3 (Singh *et al*, 2009; Teicher & Fricker, 2010), DMBA/TPA treatment induced CCL2 and CCL3 expression in WT and *Cx3cr1*^-/-^ mice to similar extents. Thus, it is likely that macrophage-derived CX3CL1 directly recruited CX3CR1-expressing M2-like macrophages to tumor tissues via using an amplifying autocrine loop.

In this carcinogenesis model, *Cx3cr1*^-/-^ mice exhibited reduced neovascularization with depressed tumor formation, compared to the case for WT mice. Indeed, CX3CR1^+^ TAMs expressed several angiogenic molecules including VEGF and MMP9, and the expression of these two molecules was reduced in *Cx3cr1*^-/-^ mice, compared to the case in WT mice. Thus, neovascularization can be ascribed at least partially to TAM-derived VEGF and MMP9. Moreover, CX3CR1 expression by CD31^+^ endothelial cells may suggest the direct involvement of the CX3CL1-CX3CR1 axis in neovascularization at tumor sites, similar to the observations in case of skin wound healing (Ishida *et al*, 2008). Nevertheless, in this skin carcinogenesis model, the CX3CL1-CX3CR1 axis could promote neovascularization directly by acting on endothelial cells and/or indirectly by inducing the production of angiogenic factors by TAMs.

Accumulating evidence indicates the crucial involvement of pro-inflammatory cytokines in DMBA/TPA-induced skin carcinogenesis. TPA-induced AP-1 activation in the epidermis is indispensable for tumor development, and requires TNFα-mediated TNF receptor signals, particularly at the early phase, in this carcinogenesis process (Moore *et al*, 1999; Arnott *et al*, 2004). IL-1 receptor-MyD88 signaling additionally contributes to keratinocyte transformation and carcinogenesis by further activating the NF-κB pathway (Cataisson *et al*, 2012). Moreover, TNFα-dependent MMP9 expression promoted epithelial cell migration during tumor promotion (Scott *et al*, 2004). Thus, these molecules can directly regulate the migration, proliferation, and transformation of keratinocytes, eventually resulting in carcinogenesis. We observed that TNFα and IL-1expression was enhanced in skin carcinogenesis of WT mice and that their expression was mainly detected in F4/80^+^ macrophages (Supplemental figure 3). Moreover, their enhanced expression was attenuated in *Cx3cr1*^-/-^ mice together with reduced F4/80^+^ macrophage recruitment, compared to the case in WT mice. Thus, locally produced CX3CL1 recruits CX3CR1^+^ macrophages, a rich source of these keratinocyte activators, thereby causing skin carcinogenesis.

Several lines of evidence have implied that CD11b^+^Gr1^+^ myeloid cells had tumor-promoting roles (Kowanetz *et al*, 2010; Qian *et al*, 2011). Di Piazza et al. have demonstrated that these cells could exert this effect by augmenting Wnt/β-catenin signaling in neighboring epithelial cells via the secretion of Wnt ligands (Di Piazza *et al*, 2012). Consistent with this observation, we also observed that TAMs were a major source of Wnt3a. Moreover, CX3CL1-CX3CR1 signaling augmented Wnt3a expression in a mouse macrophage cell line and *Cx3cr1*^-/-^ mice exhibited depressed Wnt3a expression. Furthermore, we observed enhanced COX-2 expression in F4/80^+^ macrophages. Given the capacity of prostaglandin E_2_ to trigger the Wnt/β-catenin pathway (Castellone *et al*, 2005; Shao *et al*, 2005), macrophage-derived COX-2 can activate this pathway. Nevertheless, CX3CL1 can activate the Wnt/β-catenin pathway, which is crucially involved in skin carcinogenesis, by attracting CX3CR1^+^ macrophages with a capacity to express Wnt3a and/or COX-2.

The activation of EGFR signal pathway was crucially involved in tumor proliferation including skin cancer (Tardáguila *et al*, 2013; Sibilia *et al*, 2000; Hara *et al*, 2005). In line with this, we found that the absence of CX3CR1 suppressed skin carcinogenesis with reduced TAM recruitment and less activation of EGFR signal pathway. Tardáguila and colleagues (Tardáguila *et al*, 2013) demonstrated that the CX3CL1-CX3CR1 axis promoted breast cancer through the transactivation of EGFR signals. However, CX3CL1 could not directly act on epidermal cells in chemical-induced skin carcinogenesis, since CX3CR1 was not expressed on epidermal cells. TAM produced EGF which can directly act on EGFR-expressing epidermal cells (Quail & Joyce, 2013). Thus, the attenuation of EGFR signal pathway was attributable to the reduced TAM recruitment in *Cx3cr1^-/-^* mice.

Collectively, the present observations reveal the pivotal involvement of the CX3CL1-CX3CR1 axis in several steps during chemical-induced skin carcinogenesis. Moreover, abundant CX3CL1 expression and CX3CR1^+^ macrophages in human skin cancer tissues will further support the notion that the CX3CL1-CX3CR1 axis can be a novel target for the prevention and/or treatment of human skin cancer.

## MATERIALS AND METHODS

### Reagents and antibodies (Abs)

7,12-dimethylbenz(a)anthracene (DMBA) and 12-*O*-tetradecanoylphorbol-13-acetate (TPA) were purchased from Sigma Chemical Co. (St. Louis, MO). Recombinant murine CX3CL1 was obtained from R&D Systems (Minneapolis, MN). The following monoclonal Abs (mAbs) and polyclonal Abs (pAbs) were used for immunohistochemical and immunofluorescence analyses; goat anti-mouse CX3CL1 pAbs, which cross-react with human CX3CL1, goat anti-mouse COX2 pAbs, goat anti-mouse VEGF pAbs, goat anti-mouse MMP-9 pAbs (Santa Cruz Biotechnology, Santa Cruz, CA), goat anti-human TNFα pAbs, which cross-react with mouse TNFα, goat anti-human Wnt3a pAbs, which cross-react with mouse Wnt3a, goat anti-mouse CD31 pAbs, which cross-react with human CD31, rabbit anti-mouse IL-1α pAbs (Abcam, Cambridge, UK), rabbit anti-human IL-1β pAbs, which cross-react with mouse IL-1β (Santa Cruz Biotechnology, Santa Cruz, CA), rabbit anti-human CX3CR1 pAbs, which cross-react with mouse CX3CR1 (Abnova, Walnut, CA), rat anti-mouse F4/80 mAb (clone, BM8; BMA Biomedicals, Switzerland), rabbit anti-human CD3 pAbs, which cross-react with mouse CD3 (Dako Cytomation, Kyoto, Japan), rat anti-mouse F4/80 mAb (clone, A3-1; AbD Serotec, Oxford, UK), mouse anti-human CD68 mAb (clone, 514H12; PIERCE, Aunnyvale, CA), rabbit anti-mouse Ki67 mAb (clone, D3B5; Cell Signaling, Danvers, MA), mouse anti-human HLA-DRα mAb (clone, TAL. 1B5; Dako Cytomation), mouse anti-human CD163 mAb (clone, 10D6; Leica Biosystems, Buffalo Grove, IL), rabbit anti-mouse β-catenin pAbs (Proteintech, Rosemont, IL), rabbit anti-mouse Keratin1 pAbs (BioLegend, San Diego, CA), Cy3-conjugated donkey anti-rat IgG pAbs, FITC-conjugated donkey anti-goat IgG pAbs, FITC-conjugated donkey anti-rabbit IgG pAbs (Jackson Immunoresearch Laboratories, West Grove, PA), rabbit anti-human Wnt3a pAbs which cross-reacts with mouse Wnt3a, and rabbit anti-IL-1α pAbs, which react with mouse IL-1α (Abcam, Cambridge, UK). For flow cytometric analyses, the following Abs were commercially obtained; PE-conjugated rat anti-mouse Ly-6G mAb (clone 1A8, BD Bioscience, San Jose, CA), violetFluor450-conjugated rat anti-mouse CD11b mAb (clone M1/70, TONBO, San Diego, CA), APC-conjugated rat anti-mouse F4/80 mAb (clone BM8.1, TONBO), FITC-conjugated rat anti-mouse CD3 mAb (clone 17A2, TONBO), PerCP/Cy5.5-conjugated rat anti-mouse CD45 mAb (clone 30-F11, BioLegend, San Diego, CA), FITC-conjugated rat anti-mouse CD68 mAb (clone FA-11, Bio-Rad, Oxford, UK), PE-conjugated rat anti-mouse CD206 mAb (clone MR6F3, Thermo Fisher, Waltham, MA), APC-conjugated rat anti-mouse CD86 mAb (clone GL-1, TONBO), and PerCP-conjugated goat anti-mouse CX3CR1 pAbs (R&D Systems, Minneapolis, MN). For Western blotting analyses, the following Abs were used; rabbit anti-mouse EGFR mAb (clone D38B1, #4267), rabbit anti-mouse phosphorylated (p)-EGFR mAb (clone D7A5, #3777), rabbit anti-mouse β-catenin (active) mAb (clone D13A1, #8814), and rabbit anti-mouse GAPDH mAb (clone D16H11, #5174, Cell Signaling, Danvers, MA).

### Mice

Pathogen-free 8-week-old male C57BL/6 mice were obtained from Sankyo Laboratories (Tokyo, Japan) and designated as wild-type (WT) mice. CX3CR1-deficient (*Cx3cr1*^-/-^) mice with the C57BL/6 genetic background were a generous gift from Drs. P. M. Murphy and J. L. Gao (National Institute of Allergy and Infectious Diseases, National Institutes of Health, Bethesda, MD) (Ishida *et al*, 2008). All animals were housed individually in cages under specific pathogen-free conditions during the experiments. Age- and sex-matched mice were used for the experiments. All animal experiments complied with the standards set by the Guidelines for the Care and Use of Laboratory Animals at the Wakayama Medical University.

### Human skin cancer tissues

Skin cancer tissue specimens (basal cell carcinoma (BCC), n=5, cases 1 to 5; squamous cell carcinoma (SCC), n=5, cases 6 to 10) were obtained by biopsy from the patients in the Wakayama Medical University Hospital after obtaining their informed consent for diagnosis (Supplemental Table 1). Using the densitometric tool of PhotoShop, the extents of HLA-DRα- or CD163-positivity were measured in the human BCC and SCC specimens, and were expressed as the pixel number per field (×200). The study design was approved by the Local Ethical Committee of the Wakayama Medical University Hospital.

### Skin carcinogenesis

Skin tumors were induced by two-step application of DMBA and TPA as described previous study (Wang *et al*, 2010). First, 25 µg of DMBA in 100 µl of acetone was applied onto the shaved dorsal skin of the mice. One week later, 30 µg of TPA in 100 µl of acetone was applied topically twice a week for 20 weeks. Tumor development was monitored on a weekly basis and lesions greater than 2 mm in length were counted as positive.

### Generation of bone marrow (BM) chimeric mice

The following BM chimeric mice were prepared: male *Cx3cr1*^-/-^ BM to female WT mice, male WT BM to female WT mice, male WT BM to female *Cx3cr1*^-/-^ mice, and male *Cx3cr1*^-/-^ BM to female *Cx3cr1*^-/-^ mice. BM cells were collected from the femurs of donor mice by aspiration and flushing. Recipient mice were irradiated with a radiation dose of 12 Gy using an RX-650 irradiator (Faxitron X-ray Inc., Wheeling, IL). Then, the animals intravenously received 5×10^6^ BM cells from the donor mice in a volume of 200 μl of sterile PBS under anesthesia. Thereafter, mice were housed in sterilized microisolator cages and were fed normal chow and autoclaved hyperchlorinated water for 60 days. To verify successful engraftment and reconstitution of the BM in the transplanted mice, genomic DNA was isolated from the peripheral blood and tail tissues of each chimeric mouse 30 days after BM transfer using a NucleoSpin tissue kit (Macherey-Nagel, Duren, Germany). Then, we performed PCR to detect the *Sry* gene contained in the Y chromosome (F, 5’-TTGCCTCAACAAAA-3’; R, 5’-AAACTGCTGCTTCTGCTGGT-3’). The amplified PCR products were fractionated on a 2% agarose gel and visualized by ethidium bromide staining. After durable BM engraftment was confirmed, the mice were treated with DMBA/TPA as described above.

### Histopathological and immunohistochemical analyses

At the indicated time intervals after DMBA application, skin tissues were removed, fixed in 10% formalin buffered with PBS (pH 7.2), and embedded in paraffin. Six-μm-thick sections were prepared and stained with hematoxylin and eosin. Epidermal thickness was measured using Photoshop (at 40× magnifications). Immunohistochemical analyses were also performed using anti-F4/80, anti-CD31 mAb, anti-Ki67, anti-VEGF, anti-β-catenin, or anti-Wnt3a Abs as described in a previous report (Ishida *et al*, 2012). The numbers of positive cells or CD31-positive tube-like vessels were counted on five randomly chosen visual fields at 200-fold magnifications, and the average of the five selected microscopic fields was calculated. All measurements were performed by an examiner without prior knowledge regarding the experimental procedures. A double- or triple-color immunofluorescence analysis was also conducted to identify the types of CX3CL1-, CX3CR1-, VEGF-, MMP-9-, COX-2- or Wnt3-expressing cells in the skin, as described in a previous report (Inui *et al*, 2011).

### Quantitative RT-PCR analysis

Total RNA was extracted from skin tissue using ISOGEN (Nippon Gene, Toyama, Japan), according to the manufacturer’s instructions. Next, 3 μg of total RNA was reverse transcribed to cDNA with Oligo(dT)_15_ primers using PrimeScript™ Reverse Transcriptase (Takara Bio, Shiga, Japan). The resultant cDNA was subjected to real-time PCR by using SYBR^®^ Premix Ex Taq™ II (Takara Bio) and specific primer sets (Takara Bio), as described in a previous report (Inui *et al*, 2011) (Supplemental Table 2). Amplification and detection of mRNA were conducted by using the Thermal Cycler Dice^®^ Real Time System (Takara Bio, TP800), according to the manufacturer’s instructions. To standardize the mRNA concentrations, transcript levels of β-actin were determined in parallel for each sample, and relative transcript levels were normalized based to the β-actin transcript levels.

### Flow cytometry analysis

Single-cell suspensions were prepared from wound tissue homogenates, as described in a previous report (Ishida *et al*, 2012). Contaminated red blood cells were hemolyzed using ammonium chloride solution (IMGENEX). The resulting single-cell suspensions were incubated with the antibodies for 20 minutes on ice. Isotype-matched control immunoglobulins were used to detect the nonspecific binding of immunoglobulin in the samples. The stained cells were analyzed on a CytoFLEX S system (Beckman Coulter, Brea, CA), and the obtained data were analyzed using the CytExpert 2.2 software (Beckman Coulter).

### Separation of F4/80^+^ cells by magnetic-activated cell sorting (MACS)

Skin cells were harvested, prepared into single-cell suspensions, and counted as described in a previous study (Ishida *et al*, 2012). All the incubations were conducted at 4°C for 20 min. To isolate the F4/80-positive cell population, the resultant single-cell preparation was stained with anti-F4/80 MicroBeads UltraPure mouse Abs (Miltenyi Biotec, Sunnyvale, CA). The sorting column was fixed on a MACS stand (Miltenyi Biotec) and equilibrated by using 500 μl of PBS containing 0.5% BSA. After the cell suspension was passed through a MACS MS separation column that was placed in mini MACS, the obtained F4/80^+^ cell fraction exhibited a purity of more than 95 %, as determined using a flow cytometer.

### Western blotting analysis

Skin samples were homogenized and the resultant lysates (30 μg) were electrophoresed on a 7.5% sodium dodecyl sulfate-polyacrylamide gel (SDS-PAGE) gel and transferred onto a nitrocellulose membrane. The membrane was then incubated with 1,000-fold diluted Abs against β-catenin (active), EGFR, p-EGFR, or GAPDH. After the incubation of the membrane with HRP-conjugated secondary Abs, the immune complexes were visualized using the ECL Plus System (Amersham Biosciences Corp., Piscataway, NJ), according to the manufacturer’s instructions.

### In vitro assay

Cells from the mouse macrophage cell line RAW264.7 cells were cultured in DMEM medium containing 10% fetal bovine serum, seeded at a density 1 × 10^6^ cells/well into six-well plates, and cultured overnight. After the cells were stimulated further with various concentrations of recombinant mouse CX3CL1 for 2 h at 37°C, the supernatants were collected, and the Wnt3a protein levels in these supernatants were determined using a commercially available ELISA kit (My Biosource, San Diego, CA), according to the manufacturer’s instructions. The detection limits of Wnt3a was > 23.5 ng/ml.

### Statistical analysis

Data were expressed as the mean ± SEM. For the comparison between WT and *Cx3cr1*^-/-^ mice at multiple time points, two-way ANOVA, followed by Dunnett’s post-hoc test, was used. To compare the values between two groups, unpaired Student’s *t* test was performed. In case of the series of CCL3 stimulations of RAW264.7 cells for the in vitro and the flow cytometric analysis, one-way ANOVA, followed by Dunnett’s post hoc test, was used. *P* < 0.05 was considered statistically significant. All statistical analyses were performed using the Statcel3 software under the supervision of a medical statistician.

### Study approval

Human samples were obtained under the approval by the Institutional Review Boards of Wakayama Medical University. Informed consent was received from participants prior to inclusion in the study. All animal experiments were approved by the Committee on Animal Care and Use at Wakayama Medical University. All methods were performed in accordance with the relevant guidelines and regulations.

## Acknowledgments

This work was supported in part by Grants-in-Aids for Scientific Research (A) (grant 20249040, to T. Kondo) and for Young Scientists (A) (grant 20689015, to Y. Ishida) from the Ministry of Education, Culture, Science, and Technology of Japan and by Research Grant on Priority Areas (to T. Kondo) from Wakayama Medical University, and MEXT Joint Usage/Research Center, operated by Cancer Research Institute, Kanazawa University.

## Competing interests

The authors declare no competing financial interests.

## AUTHOR CONTRIBUTIONS

Y.I and T.K formulated the hypothesis and designed the project; Y.I performed the main experiments; A.K provided technical assistance and discussion; Y.K and M.N helped with some experimental procedures; Y.Y and F.F helped to collect human skin cancer samples; N.M and T.K oversaw the experiments and provided the main funding for the project; Y.I, N.M, and T.K participated in writing the manuscript.

## Data availability

No datasets were generated or analyzed during the current study.

## The Paper Explained

### Problem

CX3CL1-CX3CR1 axis has been associated with various diseases. Previous studies have shown that CX3CL1 promotes cancer metastasis using various routes including bloodstream, lymphatic vessels, and nerves, whereas CX3CL1 prevents glioma invasion by promoting tumor cell aggregation and eventually reducing their invasiveness. The roles of CX3CL1-CX3CR1 axis in skin carcinogenesis still remains unknown.

### Results

DMBA/TPA-induced tumor incidence in *Cx3cr1^-/-^* mice was significantly reduced in comparison to that in WT mice. Infiltration of CX3CR1^+^ tumor-associated macrophages with M2-like phenotypes and the expression levels of angiogenic molecules including VEGF and MMP-9 were decreased in the skin tumor tissues of *Cx3cr1^-/-^* mice compared with WT mice. Using macrophage cell line RAW264.7 in vitro, we found that CX3CL1 associated with Wnt3a, which have tumor-promoting roles. Reduced expression of Wnt3a was identified in DMBA/TPA-induced skin tissues of *Cx3cr1^-/-^* mice compared to that in WT mice.

### Impact

These findings indicated that CX3CR1 deficiency suppressed skin carcinogenesis through the inhibition of the inflammatory tumor microenvironment by the downregulation of VEGF, MMP-9, and Wnt3a in M2-macrophages. Moreover, abundant CX3CL1 expression and CX3CR1^+^ macrophages in human skin cancer tissues further support the notion that CX3CL1-CX3CR1 axis can be a novel target for preventing and/or treating skin cancers.

## Supplemental Material

**Supplemental Figure 1.**
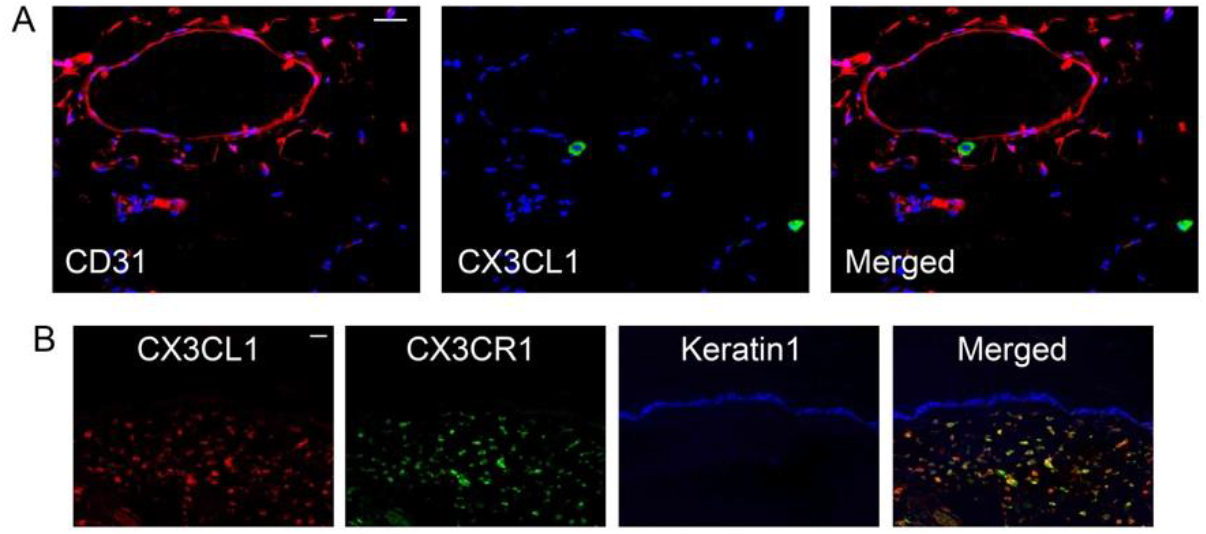
Cell types expressing CX3CL1 and CX3CR1 in the skin of DMBA/TPA-treated WT mice. (A) CD31^+^ endothelial cells did not express CX3CL1 in the skin. Scale bar, 20 μm. (B) Almost CX3CL1-expressing monocytes were also express CX3CR1 in the skin. Representative results from six individual animals are shown. Signals were merged digitally. Scale bar, 20 μm.

**Supplemental Figure 2.**
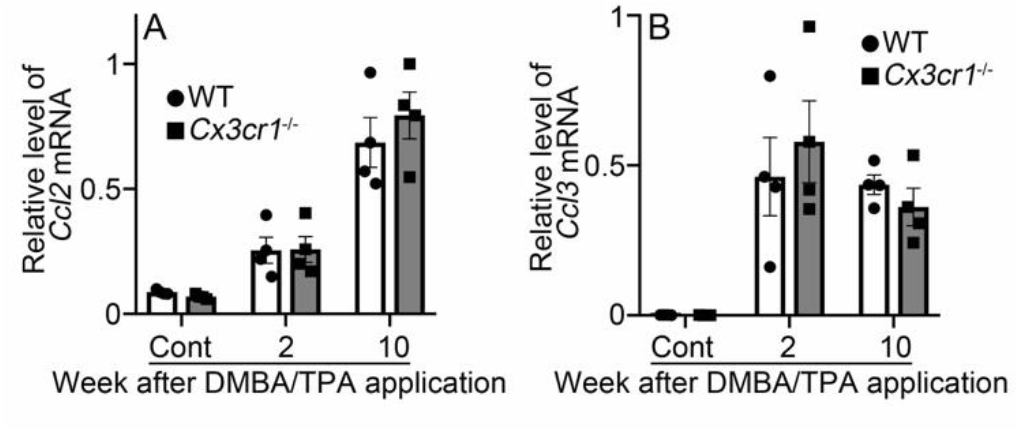
Intradermal mRNA expression of *Ccl2* (A) and *Ccl3* (B) in WT and *Cx3cr1*^-/-^ mice after DMBA/TPA treatment. Quantitative RT-PCR analyses were carried out. Values represent mean ± SEM (n=4).

**Supplemental Figure 3.**
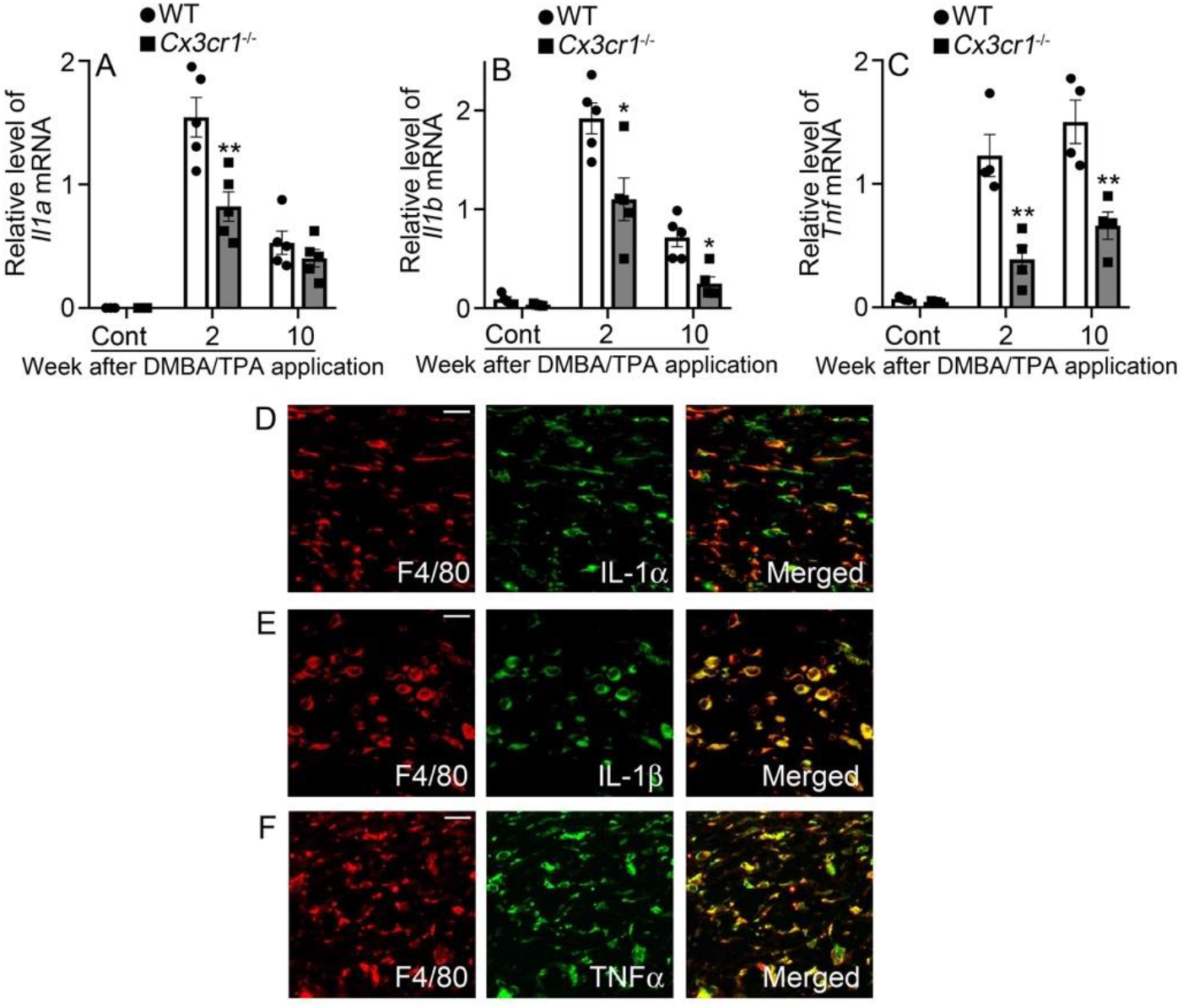
Evaluation of intradermal expression of *Il1a, Il1b* and *Tnf* in WT and *Cx3cr1*^-/-^ mice. (A-C) Intradermal gene expression of *Il1a* (A), *Ilb* (B) and *Tnf* (C) in WT and *Cx3cr1*^-/-^ mice after DMBA/TPA treatment. Values represent mean ± SEM (n=4). \**P* < 0.05, \*\**P* < 0.01, WT vs. *Cx3cr1*^-/-^ mice, by 2-way ANOVA followed by Dunnett’s post-hoc test. (D-F) Cell types expressing IL-1α (D), IL-1β (E) and TNF-α (F) in the skin of DMBA/TPA-treated WT mice. Representative results from six individual animals are shown here. Signals were merged digitally. Scale bar, 20 μm.

**Supplemental Table 1.** Profile of each patient with skin cancer

| Case No. | Type | Age | Sex | Site |
| --- | --- | --- | --- | --- |
| 1 | BCC | 86 | M | Face |
| 2 | BCC | 75 | F | Face |
| 3 | BCC | 60 | F | Face |
| 4 | BCC | 39 | F | Head |
| 5 | BCC | 59 | M | Face |
| 6 | SCC | 96 | F | Face |
| 7 | SCC | 68 | M | Hand |
| 8 | SCC | 90 | M | Finger |
| 9 | SCC | 90 | M | Face |
| 10 | SCC | 86 | M | Face |

BCC: Basal cell carcinoma, SCC: Squamous cell carcinoma

**Supplemental Table 2.**
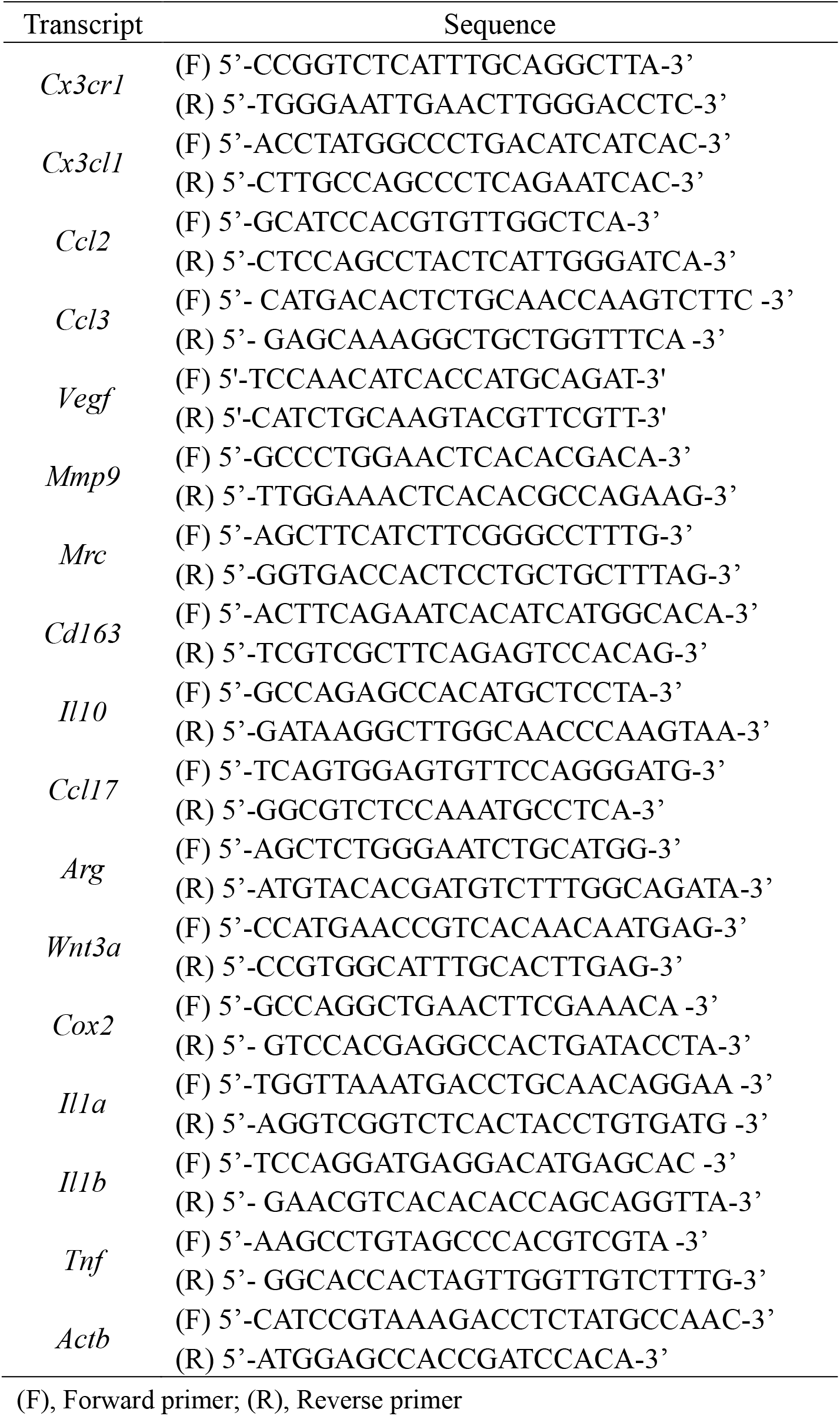
Sequences of primers used for real-time RT-PCR

## Notes

The authors declare no potential conflicts of interest.

